# Yeast Ixr1 mediates the DNA replication stress response through it HMGB DNA binding domains and interaction with checkpoint Mrc1

**DOI:** 10.1101/2023.05.30.542938

**Authors:** Siying Teng, Yi wang, Jingyuan Jiang, Mengyuan Li, Yingxin Liu, Yangying Guan, Anhui Wei, Zhongyi Cong, Xinmin Zhang

## Abstract

**Background:** High mobility group box (HMGB) family protein Ixr1 has been shown to be involved in DNA damage repair, however, its role and mechanism remain largely unclear.

**Methods:** Genes of *S. cerevisiae* were deleted or tagged with myc, GFP, or mcherry using the lithium acetate method. Sensitivity of strains to hydroxyurea (HU), methyl methanesulfonate (MMS), camptothe-cin (CPT), 4-nitroquinoline N-oxide (4-NQ), or Zeocin was tested. Distribution of GFP or mcherry fusion proteins was visualized with laser scanning confocal microscopy. RNA-seq was used to determine differential gene expression between mutant and control strains.

**Results:** Ixr1 deletion (ixr1Δ) mutant strain was sensitive to HU. Additionally, phosphorylation of effector of DNA damage checkpoint kinase Rad53 was lower in ixr1Δ than WT. Deletion of DNA damage checkpoint mediators ixr1Δ Rad9Δ was more sensitive to HU than ixr1Δ or Rad9Δ, and ixr1Δ mrc1Δ had similar sensitivity to HU as mrc1Δ but stronger than ixr1Δ. Deletion of ribonucleotide reductase inhibitors sml1Δ or crt10Δ didn’t reduce the sensitivity of ixr1Δ induced by HU. Repli-cation fork nuclease exo1Δ ixr1Δ or helicase sgs1Δ ixr1Δ double deletions were more sensitive to HU than single deletion. In addition, laser scanning confocal microscopy imaging indicated that in response to HU, Ixr1 may be in the same pathway as Mrc1, possibly downstream. Gene Ontol-ogy enrichment analysis of differentially expressed genes (DEGs) between ixr1Δ and wildtype, untreated and treated with HU, confirmed that Ixr1 plays an important role in regulating the transcription of genes related to DNA replication or DNA damage repair. We also found that, re-gardless of HU exposure, Ixr1 localized to the nucleus and may bind DNA through its two HMG-boxes.

**Conclusion:** Ixr1 participates in the DNA replication stress response through a DNA damage checkpoint pathway mediated by Mrc1, and regulates expression of genes related to DNA damage repair.

## Introduction

Genomic DNA is constantly damaged by various endogenous and exogenous agents, such as intracellular metabolites(Bist et al. 2017), reactive oxygen species(Chalissery et al. 2017), mismatches and impediments in DNA replication(Cortez 2015; Kunkel and Erie 2015), radiation(Mullenders 2018), alkylating agents(Klapacz et al. 2016), and crosslinking agents(Klages-Mundt and Li 2017). Hydroxyurea (HU), methyl methanesulfonate (MMS), Camptothecin (CPT), 4-Nitroquinoline N-oxide (4NQO), and Zeocin are genotoxins widely used in the study of DNA damage repair(Liu et al. 2022; Nogalski and Shenk 2020; Siler et al. 2017). Ribonucleotide reductase (RNR) is a key enzyme that mediates the synthesis of deoxyribonucleotides, the DNA precursors, for DNA synthesis in every living cell(Lammers and Follmann 1984). HU inhibits the activity of the RNR small subunit (R2) by affecting stability of the iron center (Nyholm et al. 1993). This results in depletion of the dNTP pool and slowing/stalling of the replication fork(Cobb et al. 2003). MMS is an alkylating agent that modifies both guanine (7-methylguanine) and adenine (3-methlyladenine) causing base mispairing and replication block, respectively(Beranek 1990). CPT, a topoisomerase I inhibitor, forms a cleavable complex with topoisomerase I and single-stranded DNA. When the complex meets a replication fork, the collision leads to irreversible double-strand breaks (DSBs)(Hsiang et al. 1985; Yamauchi et al. 2011). 4NQO, an ultraviolet light-mimetic agent, can produce chromosomal breaks or aberrations, and DNA adducts. (Darroudi et al. 1989; Felkner and Kadlubar 1968). Zeocin uses Cu^2+^ or Fe^3+^ ions as a cofactor to form single-stranded nicks which can be rapidly converted into DSBs, the predominant type of lesion detected after treatment with Zeocin(Chen et al. 2008).

To maintain genome integrity, cells may initiate the DNA damage response (DDR), activate checkpoint kinases(Lanz et al. 2019), and/or arrest the cell cycle in response to DNA damage(Lydall and Weinert 1996). DNA damage is repaired through multiple mechanisms depending on the type of DNA damage, including base excision repair (BER), nucleotide excision repair (NER), homologous recombination (HR), or nonhomologous end joining (NHEJ)(Chatterjee and Walker 2017; Santivasi and Xia 2014). If DNA damage is too extensive to be repaired, DDR will lead to apoptosis or cellular necrosis(Azzopardi et al. 2017). Error prone translesion DNA synthesis (TLS) may lead to mutagenesis, even carcinogenesis(Sasatani et al. 2020; Zhang 2020). In response to DDR, DNA damage checkpoint sensor kinase Mec1 is activated by phosphorylation and mediators Rad9 and Mrc1 transduce signals to the effector kinase, *Rad53*, to initiate downstream phosphorylation in *saccharomyces cerevisiae* (Pardo et al. 2017).

*Ixr1* codes for a high mobility group box (HMGB) family transcription factor that is involved in cisplatin sensitivity and DNA repair. Ixr1 regulates hypoxic genes during normoxia, hypoxia and oxidative stress. Cisplatin is a widely used chemotherapy drug. Cisplatin-DNA adducts stall replication forks and arrest the cell cycle at S phase(Malinge et al. 1999; Martens-de Kemp et al. 2013). Ixr1 binds DNA intrastrand cross-links formed by cisplatin through its two HMG-boxes causing a bend in the DNA that blocks excision repair of cisplatin-DNA adducts (McA’Nulty and Lippard 1996; Vizoso-Vázquez et al. 2017). Ixr1 also binds directly to the promoter of genes related to rRNA and regulates transcription and ribosome biogenesis in yeast(Vizoso-Vazquez et al. 2018). Reactive oxygen molecules from intracellular metabolism or the extracellular environment may induce DNA damage and activate the DNA damage response in cells. During transition from normoxia to hypoxia, Ixr1 regulates transcription of genes involved in aerobic repression such as *cox5b*, *tir1* and *hem13*(Castro-Prego et al. 2010). Ixr1 is also a regulatory transcription factor that binds the *rnr1* promoter region and upregulates its expression during replication stress caused by low dNTPs(Tsaponina et al. 2011). DNA damage repair is a complex process, and more functions of Ixr1 in DNA damage repair need to be found and elaborated upon.

To determine how Ixr1 responds to replication stress in yeast, we investigated its distribution and expression, its interaction with DNA damage checkpoint proteins, and its regulation of genes related to DNA damage induced by HU. The results of the current study suggest that Ixr1 is localized in the nucleus and interacts with or downstream of the mediator checkpoint Mrc1 to regulate gene expression related to DNA damage. Additionally, our data indicates that two HMG-boxes are important for Ixr1 localization to the nucleus and response to replication stress induced by HU.

## Materials and methods

### Yeast strains and primers

Genotypes of yeast strains and primers used in this study are listed in Table 1 and Table S1, respectively. All primers were synthesized by Sangon Biotech (Shanghai, China). All plasmids came from Addgene and were ordered by sinozhongyuan (Beijing, China). The *ixr1* coding region in strains 1, 2, 4, 6, 8, 10 and 12 was replaced by the KanMX4 cassette PCR-amplified with primers 1 and 2 from the pFA6a-KanMX, and verified by PCR using primers 4 and 5. Coding regions of *Rad9* (strains 2,3), *sml1* (strains 6,7), *crt10* (strains 8,9), *exo1* (strains 10,11) and *sgs1* (strains 12,13) were replaced by the Hygromycin B resistance gene cassette PCR-amplified with primers 6/7 (*Rad9)*, 18/19(*sml1*), 22/23 (*crt10*), 26/27 (*exo1*), and 30/31 (*sgs1*) from the pFA6a-hphMX6 and verified by PCR using primers 9/11 (*Rad9*), 20/21 (*sml1*), 24/25 (*crt10*), 28/29 (*exo1*), and 32/33 (*sgs1*), respectively. Coding regions of *mrc1*(strains 4,5) were replaced by the clonNAT resistance gene cassette PCR-amplified with primers 12 and 13 from the pFA6a-natMX and verified by PCR using primers 16/17. Ixr1was tagged with green fluorescent protein (GFP) (strains 14, 17, 18, 19) by inserting the sequence of GFP-KanMX before the stop codon of *ixr1* by PCR-amplified with primers 2/3 from the pFA6a-GFP-KanMX, and verified by PCR using primers 4/5. The *Ixr1*_HMGBΔ (strain 16) was made by transforming sequence amplified with primers 2/30 from the pFA6a-KanMX and verified by PCR using primers 4/5. The *Ixr1*_HMGBΔ_GFP (strain15) was made by transforming sequence amplified with primers 2/31 from the pFA6a-GFP-KanMX and verified by PCR using primers 4/5. Rad9, Mrc1 and Rad53 were tagged with mcherry (strains 17, 18, 19) by inserting the sequence of mcherry-clonNAT before the stop codon of *ixr1* by PCR-amplification with primers 7/8 (Rad9), 12/14 (Mrc1), 36/37 (Rad53) from the pFA6a-mcherry-clonNAT and verified by PCR using primers 10/11 (Rad9), 15/16 (Mrc1), and 38/39 (Rad53).

### DNA damage assay

Single colonies picked from plates were inoculated and cultured in YPD (1% yeast extract (LP0021B, Oxoid, Thermo Scientific), 2% peptone (LP0037B, Oxoid, Thermo Scientific), and 2% glucose (G116303, Aladdin, Shanghai, China)) liquid medium at 30℃ overnight. Cultures were adjusted to an OD600 of approximately 1.0, 10x serial diluted, and spotted on YPD plates (YPD+2% agar) with or without MMS (M301750, Aladdin, Shanghai, China), HU (H106352, Aladdin, Shanghai, China), CPT (B21343, Yuanye, Shanghai, China), or Zeocin(ant-zn-05, Invivogen). Plates were incubated for 2 days and scanned by Epson V600 scanner.

### Cell cycle analysis

To study the effect of genotoxins on cell cycle, cultures in log phase (OD600 ∼0.3) were arrested at G1 phase by adding α-factor twice (HY-P1482, MedChemExpress, Shanghai, China) (final concentration 1µg/ml) for 1.5 h each. Cells were collected and washed with cold water twice and then cultured with prewarmed fresh YPD with or without genotoxins. Cells were then collected at multiple time points (0, 15, 30, 45 and 60 min). The collected cells were washed with cold water and fixed in 70% ethanol overnight at 4°C. Cells were washed with 50 mM NaCitrate (S11110, Yuanye, Shanghai, China) (pH = 7.4), resuspend in 50 mM NaCitrate containing 0.25 mg/ml of boiled RNase A (2158, Takara, Kusatsu, Japan), at 37℃ for 2 h, and treated with Proteinase K (A004240, Sangon Biotech, Shanghai, China) for 20 min at 50 ℃. Next, 50 mM NaCitrate containing 20 µg/ml of propidium iodide was added and cells were stored at 4 ℃ overnight. Cells were analyzed on a FACSCalibur (BD Biosciences, San Jose, CA, USA). Data acquisition was analyzed using FlowJo software (BD Biosciences, San Jose, CA, USA).

### RNA-seq and data analysis

RNA purification, library preparation, and RNA-sequencing were performed by Novogene (Beijing, China). Raw RNA-seq data was submitted to the Sequence Read Archive (SRA) of National Center for Biotechnology Information. BioProject ID xx. Data were analyzed using R software (http://www.r-project.org). Read quality was assessed and adapter sequences removed using Rfastp (Chen et al. 2018). Clean reads were mapped to the *s. cerevisiae* reference genome (assembly R64) from NCBI using Rsubread(Liao et al. 2019). Differentially expressed genes (DEGs) were analyzed using EdgeR(Robinson et al. 2010). Gene Ontology (GO) and Kyoto Encyclopedia of Genes and Genomes (KEGG) enrichment analysis were performed and visualized using clusterProfiler (Wu et al. 2021). Heatmaps were constructed using pheatmap(Kolde 2019).

### Western blot

In order to determine the effect of HU on expression of Ixr1, 3HA epitopes were fused to the C-terminal of Ixr1. Exponentially growing yeast cells were treated or untreated with 150 mM HU for 30min. To detect the effect of Ixr1 on the phosphorylation of Rad53, exponentially growing W303 and *ixr1*Δ strains were synchronized at G1 phase by α-factor, then released, and treated with 150 mM HU for 30 min. Whole cell extracts were made by glass beads beating with bead beater (Ixr1) (DeCaprio and Kohl 2020) or TCA precipitation (Rad53)(Szymanski and Kerscher 2013). Protein concentration was quantified with the BCA Protein Assay Kit (P1002) from Beyotime (Shanghai, China). Proteins were separated by SDS-PAGE followed by transfer to PVDF membrane (88518, ThermoFisher) using the Trans-Blot Turbo Transfer System (JUNYI, Beijing, China). HA rabbit polyclonal antibody (AP0005, 1:2000) was purchased from Bioworld Technology (Nanjing, China), Rad53 rabbit polyclonal antibody (ab104232, 1:2500) was purchased from Abcam (Cambridge, England). β-Actin mouse monoclonal antibody (AF5001,1:1000), secondary antibody HRP-labeled goat anti-rabbit IgG(H+L) (A0208,1:10000), secondary antibody HRP-labeled goat anti-mouse IgG(H+L) (A0216,1:5000) and ECL reagent (P0018FS) were purchased from Beyotime (Shanghai, China).

## Results

### Ixr1 is involved in HU-induced replication stress response

To examine the effect of Ixr1 on cellular resistance to different genotoxins, growth phenotypes of WT and *ixr1*Δ strains on YPD plates containing HU, MMS, 4NQO, CPT, or Zeocin were examined. *Ixr1* deletion decreased resistance to HU but no obvious growth pattern changes were observed on the plates containing MMS, 4NQO, CPT, or Zeocin (Fig. 1A). Western blot and flow cytometry showed that the expression of Ixr1 was upregulated following HU treatment (Fig. 1B, 1C). In yeast, DNA damage checkpoint signaling cascades are activated from detectors (Mec1 and Tel1) to effector (Rad53) in response to DNA damage. Phosphorylation of Rad53 is a marker of activation and related to cell cycle arrest, origin collapse suppression, and dNTP upregulation(Lanz et al. 2019; Moriel-Carretero et al. 2019; Pardo et al. 2017). Rad53 was a slightly lower in *ixr1*Δ mutants exposed to HU in comparison to WT (Fig. 1D). Ixr1 is a transcription factor involved in regulation of hypoxic genes expression by binding regulatory sequences, as demonstrated by electrophoretic mobility shift assay(Vizoso-Vázquez et al. 2017). To verify whether the distribution of Ixr1 in cells differs under HU induced replication stress, GFP was fused to the C-terminal of Ixr1. The results showed that Ixr1 is localized to the nucleus with or without HU treatment (Fig. 1E). HU depletes the cellular pool of dNTPs, the raw material for DNA synthesis, by inhibiting the activity of RNRs(Pardo et al. 2017; Poli et al. 2012). DNA is synthesized in the S phase of the cell cycle. We analyzed the effect of Ixr1 deletion on the progression of S phase. Flow cytometry results indicated Ixr1 deletion has no effect on the progression of S phase in HU treated cells (Fig. 1F).

**Fig 1.**
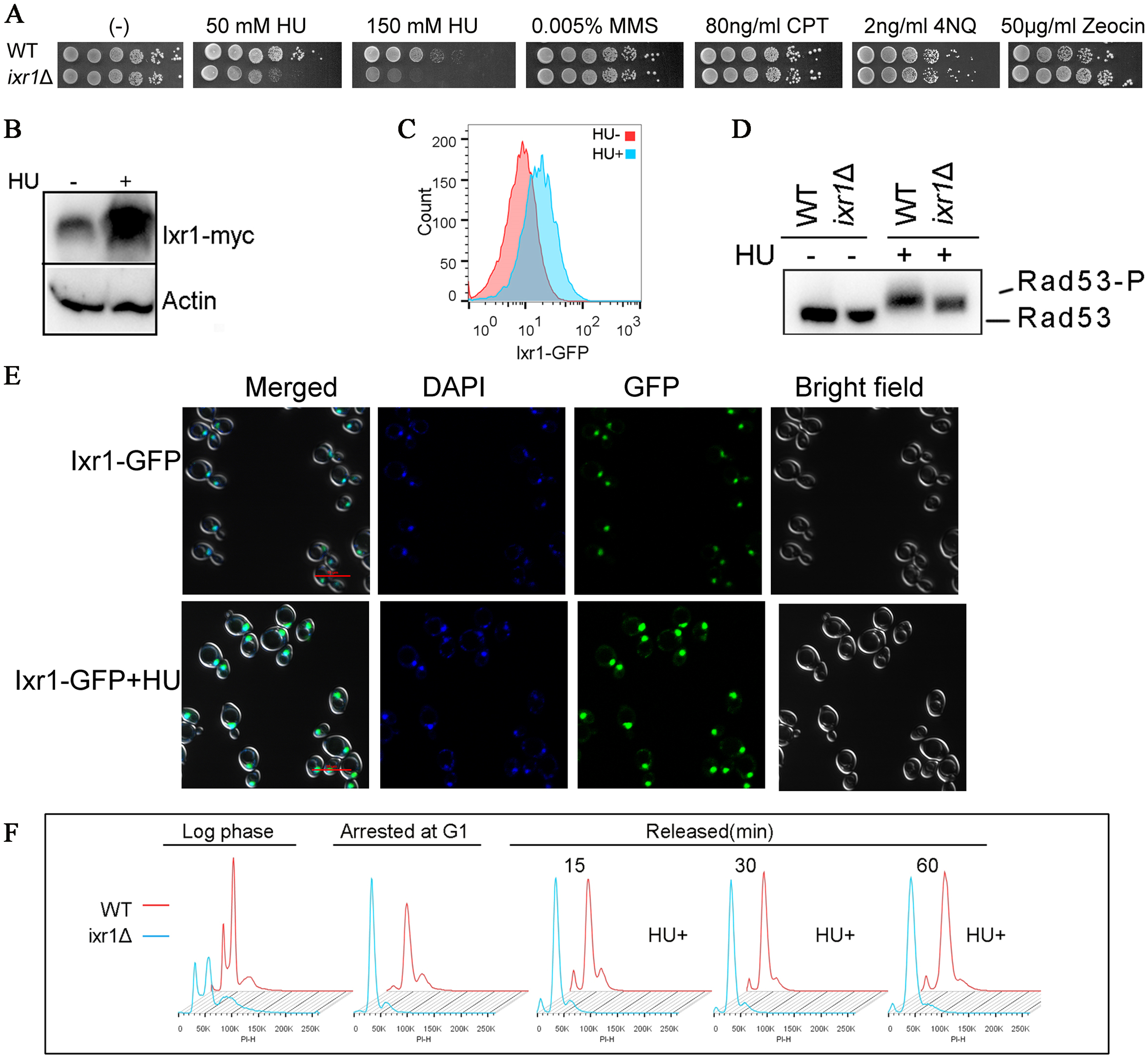
Ixr1 is involved in HU-induced replication stress response. A. Growth phenotype of wildtype W303 and *ixr1*Δ strains on YPD plates or YPD plates containing HU, MMS, 4NQO, CPT or Zeocin. B. Expression of Ixr1 detected by western blot. Ixr1 fused with myc was detected using myc antibody. Actin was used as loading control. C. Expression of Ixr1 detected by flow cytometry. (x-axis represents intensity of Ixr1-GFP signal; y-axis represents number of cells). D. Phosphorylation of Rad53 detected by western blot. (- represents yeast cells without HU treatment, + represents cells with HU treatment, Rad53-P represents phosphorylated Rad53). Variable Rad53-P migration corresponds to multiple Rad53 phosphorylation sites. E. Distribution Ixr1-GFP treated with or without HU observed by confocal laser scanning microscope. (green=GFP, blue=DAPI). F. The trend of S-phase DNA content changes over time is the same in *ixr1*Δ and WT treated with HU. (x-axis represents propidium iodide signal; y-axis represents number of cells)

### Ixr1 might act downstream of or directly with checkpoint mediator Mrc1

In yeast, DNA damage signals are transmitted from sensor Mec1 to effector Rad53 through mediators Rad9 and Mrc1(Alcasabas et al. 2001; Pardo et al. 2017). Rad9 responds to DNA damage in all stages of the cell cycle while Mrc1 is mainly involved in repairing DNA damage during S phase(Pardo et al. 2017). Replication fork arrest is induced by HU(Poli et al. 2012). Uncoupling of DNA polymerase and helicase activities causes single stranded DNA (ssDNA) gaps to form at the replication fork which go on to generate double stranded DNA breaks(Pardo et al. 2017; Sogo et al. 2002). Thus, HU-induced DNA damage includes replication stress, ssDNA breaks, and DSBs. Rad9 is a DNA damage-dependent checkpoint adaptor, transmitting checkpoint signal from sensor proteins Mec1/Tel1 to effector proteins Rad53 and Chk1p in response to DNA damage(Blankley and Lydall 2004; Sweeney et al. 2005). Since Rad53 phosphorylation was slightly lower in *ixr1*Δ cells exposed to HU than WT (Fig. 1C), we further investigated the interactions of Ixr1 that may affect the activation of Rad53 through checkpoint mediators Mrc1 or Rad9. The results showed that *Rad9*Δ *ixr1*Δ double mutants were less resistant to HU than *Rad9*Δ or *ixr1*Δ single mutants (Fig.2A). Furthermore, *mrc1*Δ *ixr1*Δ double mutants had similar resistance to HU as *mrc1*Δ but less resistance than *ixr1*Δ alone (Fig.2B). In addition, confocal laser scanning microscopy showed aggregation of Ixr1, Rad9, Mrc1 and Rad53 in response to HU induced replication stress. Ixr1, Mrc1, and Rad53 foci were distributed in the same location, while Rad9 did not co-localize with Ixr1 in most yeast cells, although it overlapped with Ixr1 in some clones. These results indicated that Ixr1 might be involved in HU induced DNA damage repair through direct interaction with or downstream of Mrc1 though not through the same signaling pathway as Rad9.

**Fig. 2.**
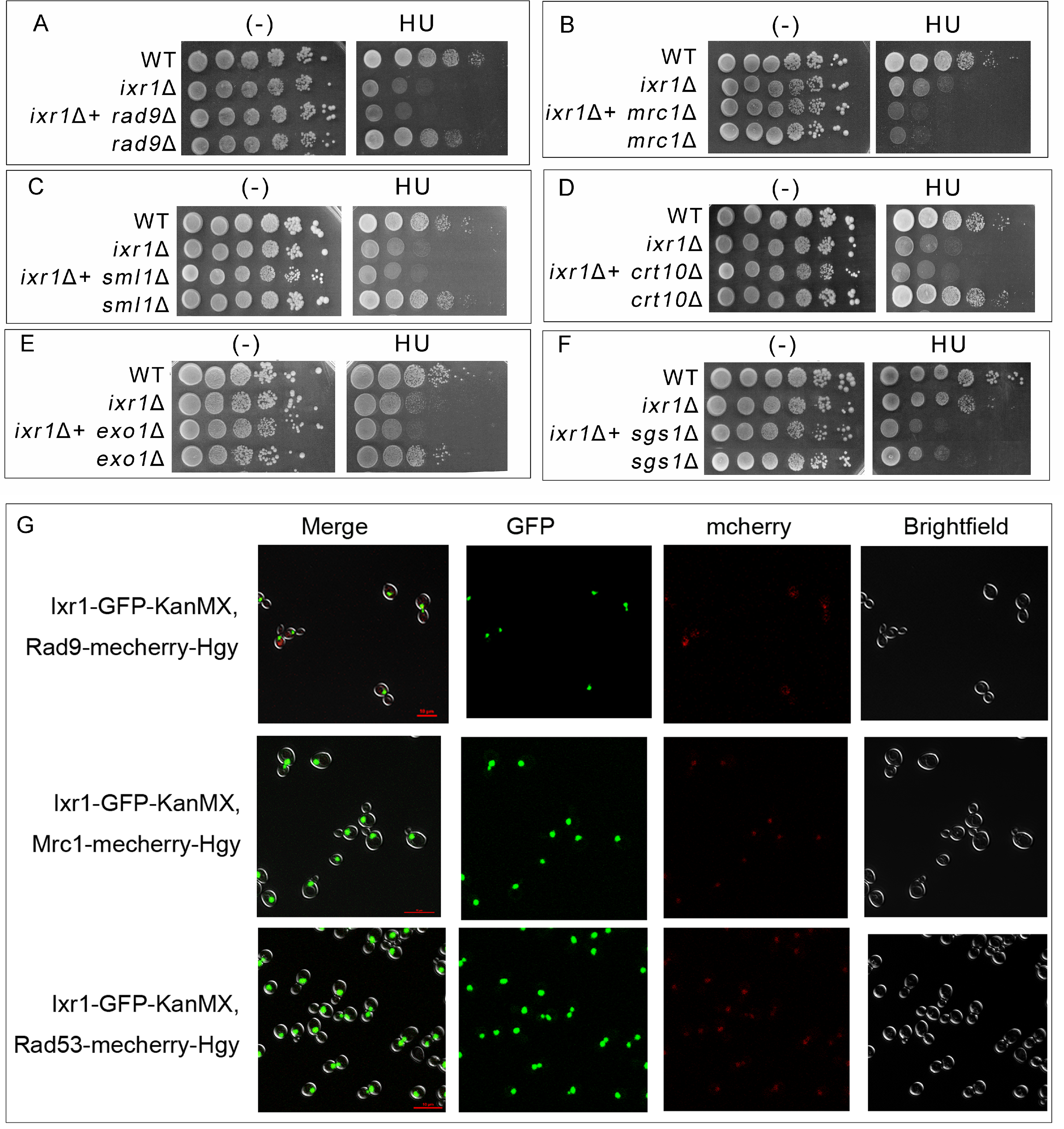
Ixr1 might act downstream of or directly with checkpoint mediator Mrc1. A-F. Growth phenotypes of *ixr1*Δ crossed with *rad9*Δ, *mrc1*Δ, *sml1*Δ, *crt10*Δ, *exo1*Δ and *sgs1*Δ in response to HU. (-) Represents the absence of HU in the medium, and HU represents the addition of a final concentration of 150 mM HU in the medium. G. Distribution Ixr1-GFP and checkpoint Rad9-mcherry, Mrc1-mcherry, Rad53-mcherry treated with 150mM HU was observed by confocal laser scanning microscope. (Green=GFP; red=mcherry).

### RNR inhibitors (Sml1 or Crt10) cannot rescue or reduce the sensitivity of *ixr1***Δ** to HU

The appropriate concentration of dNTPs is vital for cell cycle progress and DNA damage repair(Poli et al. 2012). Ribonucleotide reductases (RNRs), catalyzes the conversion of ribonucleotides to deoxyribonucleotides, which is a rate-limiting step in dNTP synthesis(Ge et al. 2001). Sml1 and Crt10 are inhibitors of RNR enzymes(Fu and Xiao 2006; Misko et al. 2016; Zhao et al. 2000). Our results showed that *sml1*Δ or *crt10*Δ in the *ixr1*Δ strain have similar growth patterns with *ixr1*Δ strain treated with HU, indicating that deletion of the two RNR inhibitors could not rescue or reduce the sensitivity to HU caused by deletion of *Ixr1*(Fig2.C-D).

### *ixr1*Δ synergistically enhanced the sensitivity of *exo1*Δ and *sgs1*Δ to HU

Exo1 plays an important role in dissociation of the replication fork replication complex and suppressing the formation of reverse replication forks during DNA replication stress(Cotta-Ramusino et al. 2005; Sogo et al. 2002). Sgs1 helps stabilize DNA polymerase at replication forks during HU treatment, recruits Rad53 kinase to stalled forks, and cooperates with Mec1 to fully activate Rad53(Cobb et al. 2003; Hegnauer et al. 2012). Immunofluorescent staining confirmed that Ixr1 is ubiquitously bound to genomic DNA (Fig. 1D). To determine if Ixr1 interacts with Exo1 and Sgs1, we analyzed the growth phenotype in *ixr1*Δ, *exo1*Δ, *sgs1*Δ single and double mutants exposed to HU induced DNA replication stress. The growth phenotypes showed that *ixr1*Δ*exo1*Δ and *ixr1*Δ*sgs1*Δ double mutants exhibit more sensitivity to HU than single mutant (Fig.2E, 2F), indicating that Ixr1 may be in different signaling pathways with Sgs1 or Exo1.

### The HMGB DNA binding domain is required for Ixr1’s functional response to HU

Ixr1 regulates multiple genes expressed under hypoxic conditions and in response to cisplatin treatment by binding promoter regions(Barreiro-Alonso et al. 2018; Castro-Prego et al. 2009; Castro-Prego et al. 2010; McA’Nulty et al. 1996). Laser scanning confocal microscopy showed that Ixr1 localizes to the nucleus in the presence or absence of HU (Fig. 1D). Ixr1 binds DNA through its two HMG-boxes (Vizoso-Vázquez et al. 2017). To demonstrate the necessity of the HMGB-box during response to HU induced replication stress, we analyzed the effect of deleting the HMGB domain (*Ixr1_HMGB_*Δ) on intracellular Ixr1 localization and sensitivity to HU in yeast. Our results showed that, under HU treatment, *Ixr1_HMGB_*Δ mutants had similar growth patterns to *ixr1*Δ mutants (Fig. 3A). Laser scanning confocal microscopy showed that Ixr1_HMGB_Δ was distributed throughout the cytoplasm but not in the nucleus in both the presence and absence of HU. These results indicate that the HMGB domain is important for Ixr1 nuclear localization as well as the response to HU treatment (Fig. 3B).

**Fig. 3.**
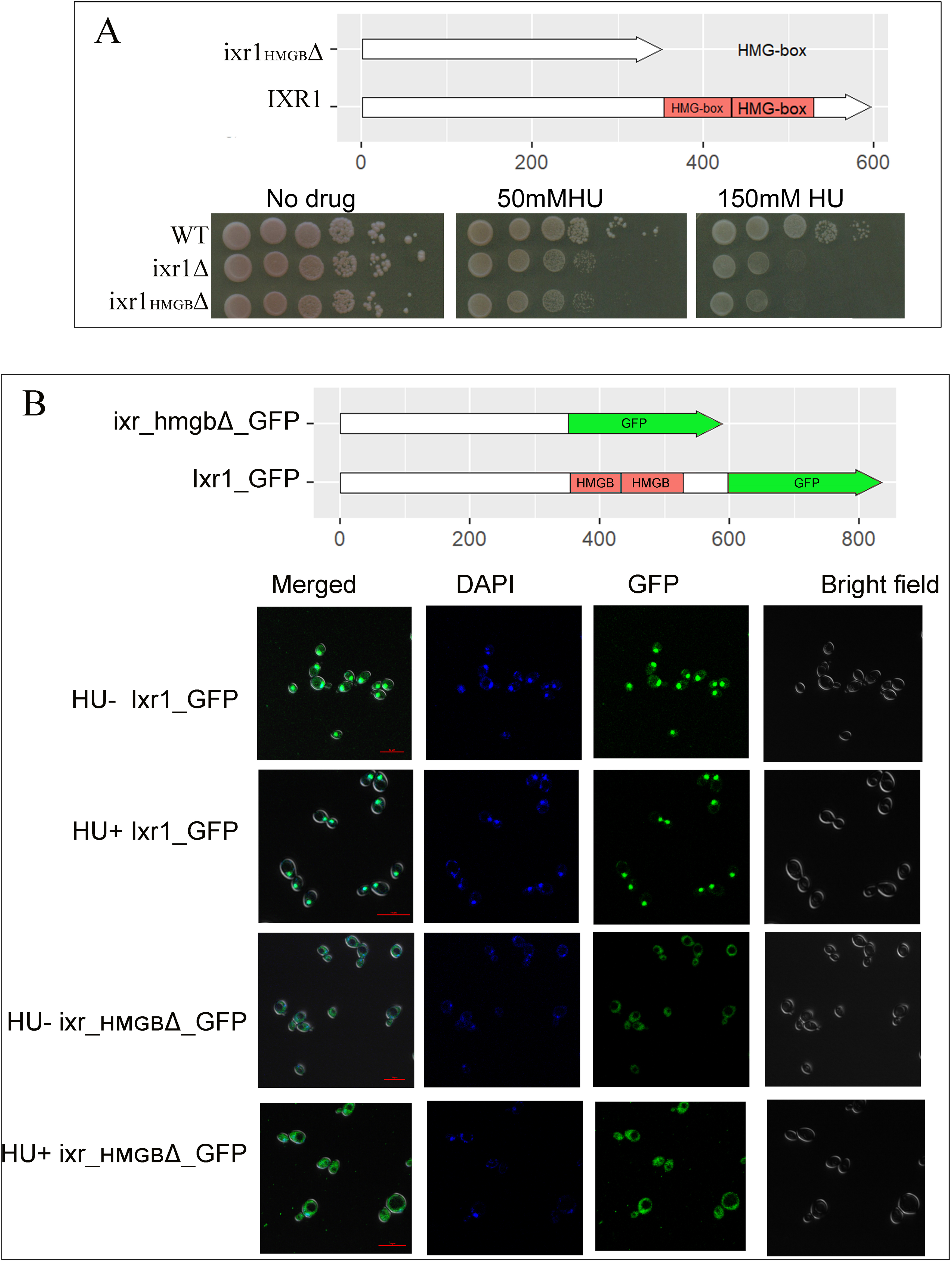
The HMGB domain is important for Ixr1 nuclear localization and the response to HU. A. The DNA binding HMGB domain is important for Ixr1’s function in response to HU. Top, a schematic of the structure of Ixr1. HMGB domain is shown as red filled boxes. Bottom, growth phenotypes of strains. WT=Ixr1 with intact structure, *ixr1*Δ=Ixr1 deleted; *ixr1__HMGB_*Δ=HMGB domain deleted from Ixr1. B. HMGB domain is important for Ixr1 aggregation in the nucleus. Top, a schematic of the structure of Ixr1 and *ixr1__HMGB_*Δ fused with GFP. HMGB domain is shown as red filled boxes and GFP is shown as green filled arrows. Bottom, distribution Ixr1-GFP and ixr1__HMGB_Δ-GFP treated with or without HU observed by confocal laser scanning microscope. (Green=GFP; blue=DAPI).

### Ixr1 regulates gene transcription related to DNA replication and damage repair

Our data shows that transcription factor Ixr1 localizes to the nucleus in yeast treated with or without HU (Fig. 1D). Logically, Ixr1 might responds to HU through regulation of gene transcription. Inhibition of ribonucleotide reductase, the rate-limiting enzyme in dNTP synthesis, by HU causes replication stress during S-phase(Pardo et al. 2017). Here, using RNA-seq, we analyzed transcriptomic changes in S-phase cells (released from α-factor arrested G1 phase into S phase after 30min) in response to HU (150mM) treatment. Genes with a fold change greater than 2X (|log2 fold change|>1) and *p*-values less than 0.05 were regarded as differentially expressed. Compared to untreated WT cells, HU treated WT cells had 1342 downregulated and 1410 upregulated genes (Fig. 4A1). Compared to WT cells, *ixr1*Δ cells had 111 downregulated and 159 upregulated genes (Fig. 4A2). Compared with untreated *ixr1*Δ cells, *ixr1*Δ HU treated cells had 925 downregulated and 627 upregulated genes (Fig. 4A3). Gene Ontology (GO) enrichment analysis in HU treated WT vs. WT and *ixr1*Δ vs. WT showed that DEGs were enriched in many DNA damage repair related biological processes, cellular components, and molecular functions, including nucleotide metabolic process, cell cycle, and cellular response to DNA damage stimulus (Fig. 4B1-2). None of these processes, however, were enriched for *ixr1*Δ HU treated vs. *ixr1*Δ untreated (Fig. 4B3). These results confirm that Ixr1 plays an important in regulating gene transcription related to DNA replication and DNA damage repair.

**Fig. 4.**
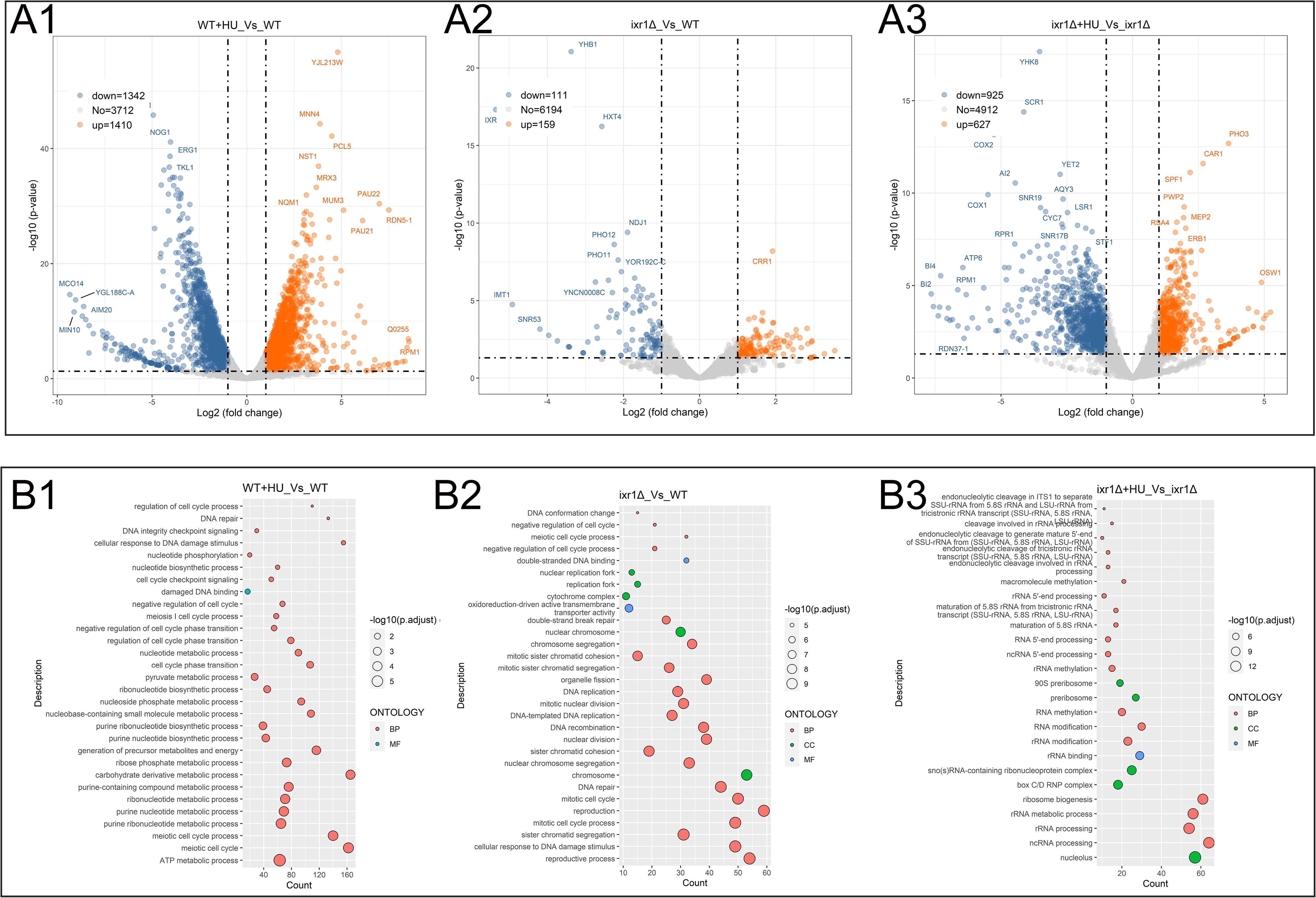
Effect of *ixr1*Δ on gene transcription in the presence and absence of HU. A. Volcano plots of differential gene expression. B. Dot-plot of enrichment analysis (based on GO) analysis of gene expression of yeast cells treated and not treated with HU.

## Discussion

Ixr1 is a member of the HMGB protein family which also includes Hmo1, Nhp10, Nhp6, and Rox1. Many HMGB proteins participate in different types of DNA damage repair through different mechanisms. In response to DNA double-strand breaks (DSBs), expression of Hmo1 decreases. Hmo1 promotes non-homologous end-joining repair by controlling DNA end resection, thus limiting error-prone alternative-end joining repair of DSBs(Panday et al. 2017a; Panday et al. 2017b). NHP10 helps recruit the Ino80 complex to DSBs(Morrison et al. 2004). Nhp6 deletion reduces histone occupancy and increases chromatin movement, decompaction, and ectopic homology-directed repair(Cheblal et al. 2020).

In this study, we found that the expression of Ixr1 is up-regulated under HU treatment. Furthermore, the *ixr1*Δ strain is sensitive to DNA damage reagent HU, but not MMS, CPT, Zeocin, or 4NQ. These results suggest that Ixr1 does not participate in the response to DNA damage caused by base modification, single strand break, or UV radiation, but rather DNA replication stress induced by HU. HU inhibits the R2 subunit of ribonucleotide reductases (RNR)(Nyholm et al. 1993). RNR is the rate-limiting enzyme in the biosynthesis of dNTPs which are essential for DNA replication and damage repair(Poli et al. 2012). Insufficient DNA replication material induced by HU leads to uncoupling of DNA helicase and replicase(Knighton et al. 2019; Pardo et al. 2017; Poli et al. 2012). The rate of DNA synthesis by DNA polymerase is consequently uncoupled with the speed of MCM2-7 helicase complex unwinding, resulting in excessive deleterious ssDNA stalling at the replication fork(Feng et al. 2006; Sabatinos and Forsburg 2015). This further induces DNA double strand breaks (DSBs). The Mre11-Rad50-Xrs2 complex recognizes DSBs and 5’-ends are nicked by Sae2. Long-range resection by exonuclease Exo1 and the Sgs1-Dna2 complex form a 3’-ssDNA, which is bound by the RPA complex. With the help of Rad52, Rad51 replaces RPA to form the Rad51 filament that seeks out and invades the homologous template(Bhat and Cortez 2018; Cejka et al. 2010; Kantake et al. 2003). DNA is then synthesized according to the template. Here, we found that deletion of *ixr1* in *exo1*Δ and *sgs1*Δ mutants synergistically increases HU sensitivity, further indicating that Ixr1 might not participate in the DSB of HR pathway.

Using western blot, Tsaponina et. Al. showed that, in response to DNA damage induced by genotoxins HU and 4-NQO, Ixr1 maintains dNTP pools by increasing the expression of RNR1(Tsaponina et al. 2011). HU inhibits the activity of RNRs, resulting in decreased dNTP concentrations(Poli et al. 2012). Sml1 and Crt10 are transcription inhibitors of RNRs(Fu and Xiao 2006; Misko et al. 2016; Zhao et al. 2001; Zhao et al. 2000). We found that deletion of *sml1* and *crt10* has no obvious recovery effect on the *ixr1*Δ phenotype induced by HU. Thus, Ixr1 might directly act on RNRs rather than the RNR inhibitors during DNA replication stress induced by HU.

During DNA replication stress induced by HU, RPA coated ssDNA triggers the checkpoint (Tel1/Mec1→Mrc1/Rad9→Rad53) response resulting in cell cycle arrest and repression of late firing replication origins as well as upregulation of dNTP synthesis to help complete replication(Gan et al. 2017; Kim and Brill 2003; Pardo et al. 2017). Mediators Rad9 and Mrc1 are checkpoint adaptor proteins that transmit signals from sensor proteins Mec1 and Tel1 to effector protein Rad53 through phosphorylation(Pardo et al. 2017). Rad9 is recruited to chromatin through interactions with histone H2A which is phosphorylated on serine 129 by the kinase Tel1 and histone H3 which is methylated on lysine 79 by histone methyltransferase Dot1(Conde et al. 2009; Toh et al. 2006; Wysocki et al. 2005). Once bound to chromatin, Rad9 is phosphorylated by Mec1 and can then recruits Rad53. Rad53 is phosphorylated by Mec1 and then autophosphorylates in a manner dependent on the adaptor Rad9, leading to full activation of Rad53 and amplification of the checkpoint signal(Gilbert et al. 2001; Schwartz et al. 2002; Sweeney et al. 2005). Mrc1 not only plays a vital role in DNA replication under normal conditions, but is also a mediator for Rad53 activation under replication stress. Mrc1 modulates origin firing, replication fork assembly, stability, and progress during cell cycle progression (Baretic et al. 2020; Randell et al. 2010). In the presence of HU, Mrc1 cooperates with Hsk1 to activate late/dormant origins and cooperates with Ctf18, Tof1, MCM-Cdc45 helicase and other factors to protect replication forks(Calzada et al. 2005; Crabbé et al. 2010; Matsumoto et al. 2017). In this study, we show that phosphorylation of Rad53 is slightly lower in *ixr1*Δ mutants than in WT strains exposed to HU. *Ixr1* deletion enhances the sensitivity of *Rad9* deletion mutants to HU but does not change the sensitivity of *mrc1* deletion mutants. Additionally, *mrc1*Δ mutants are more sensitive than *ixr1*Δ mutants which suggests that Ixr1 may act downstream or interact directly with Mrc1.

Ixr1 is reported to be a transcriptional factor that regulates the expression of hypoxic and aerobic genes by binding their promoter regions during normoxia and hypoxia (Barreiro-Alonso et al. 2018; Castro-Prego et al. 2009; Castro-Prego et al. 2010; Vizoso-Vázquez et al. 2012). Ixr1 also binds DNA intrastrand cross-links formed by cisplatin (McA’Nulty et al. 1996). *Hem13* transcriptional activation is dependent on two tandem HMGB domains of Ixr1 (Vizoso-Vázquez et al. 2017). In this study, we found that Ixr1 localizes to the nucleus with or without HU treatment. Conversely, Ixr1 lacking HMGB domains is distributed throughout the cell but not in the nucleus. GO enrichment analysis of DEGs indicated that Ixr1 plays an important role in regulating the transcription of genes related to DNA replication and damage repair, such as, cellular response to DNA damage stimulus, chromosome, DNA-templated DNA replication. We therefore hypothesize that Ixr1 regulates gene transcription by binding chromatin through its two HMG boxes.

## Conclusion

Ixr1 participates in the response to DNA replication stress induced by HU. It may act downstream of or directly with checkpoint Mrc1 and regulate gene transcription by binding chromatin through its two HMG boxes.

## Acknowledgement

The authors would like to thank the science and technology development project of Jilin Province (grant No.20220402043GH).

## Supplementary Materials

Table S1: Sequences of primers used in this study.

## Data Availability Statement

The raw data of RNA-seq were submitted to the Sequence Read Archive (SRA) of National Center for Biotechnology Information. BioProject ID PRJNA973186.

---

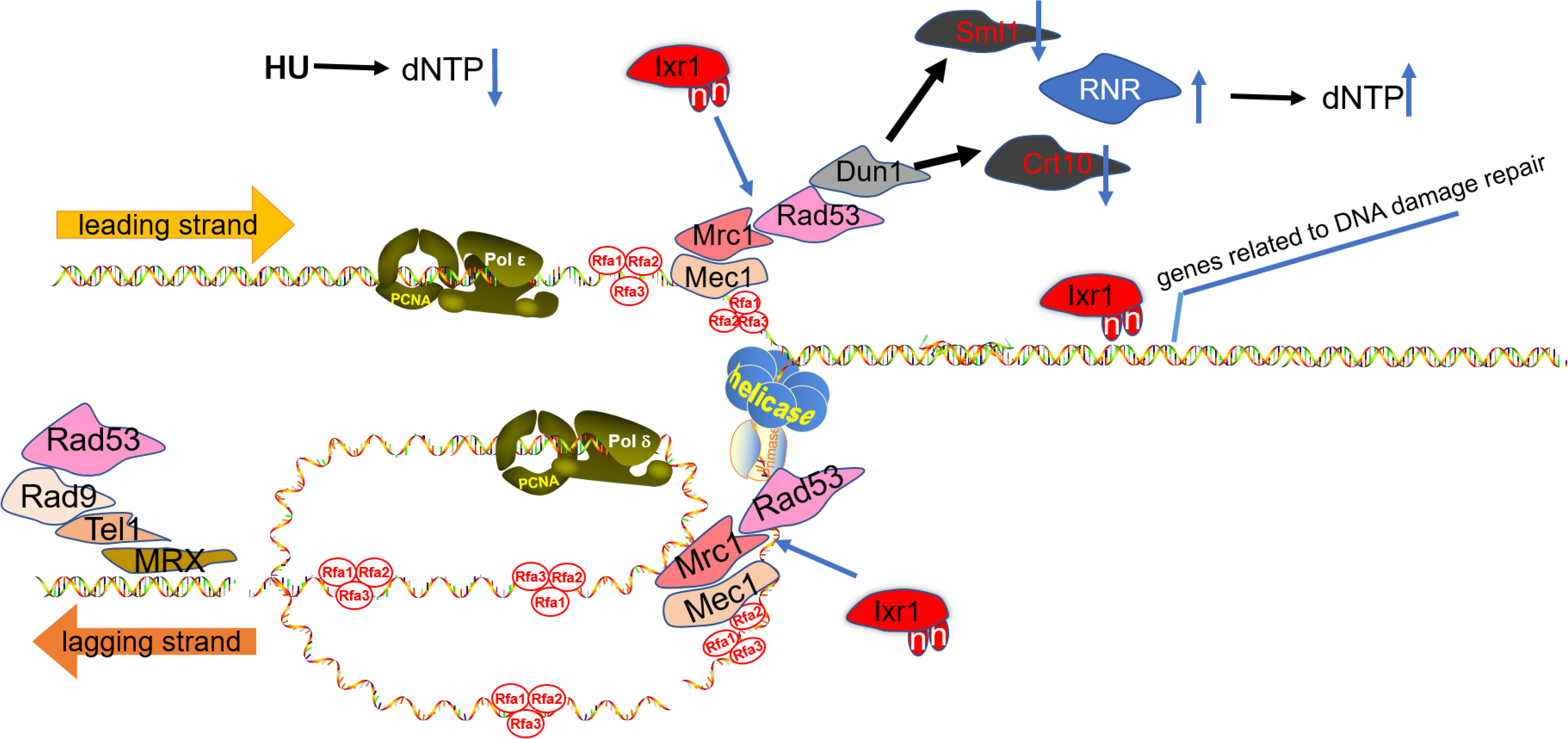

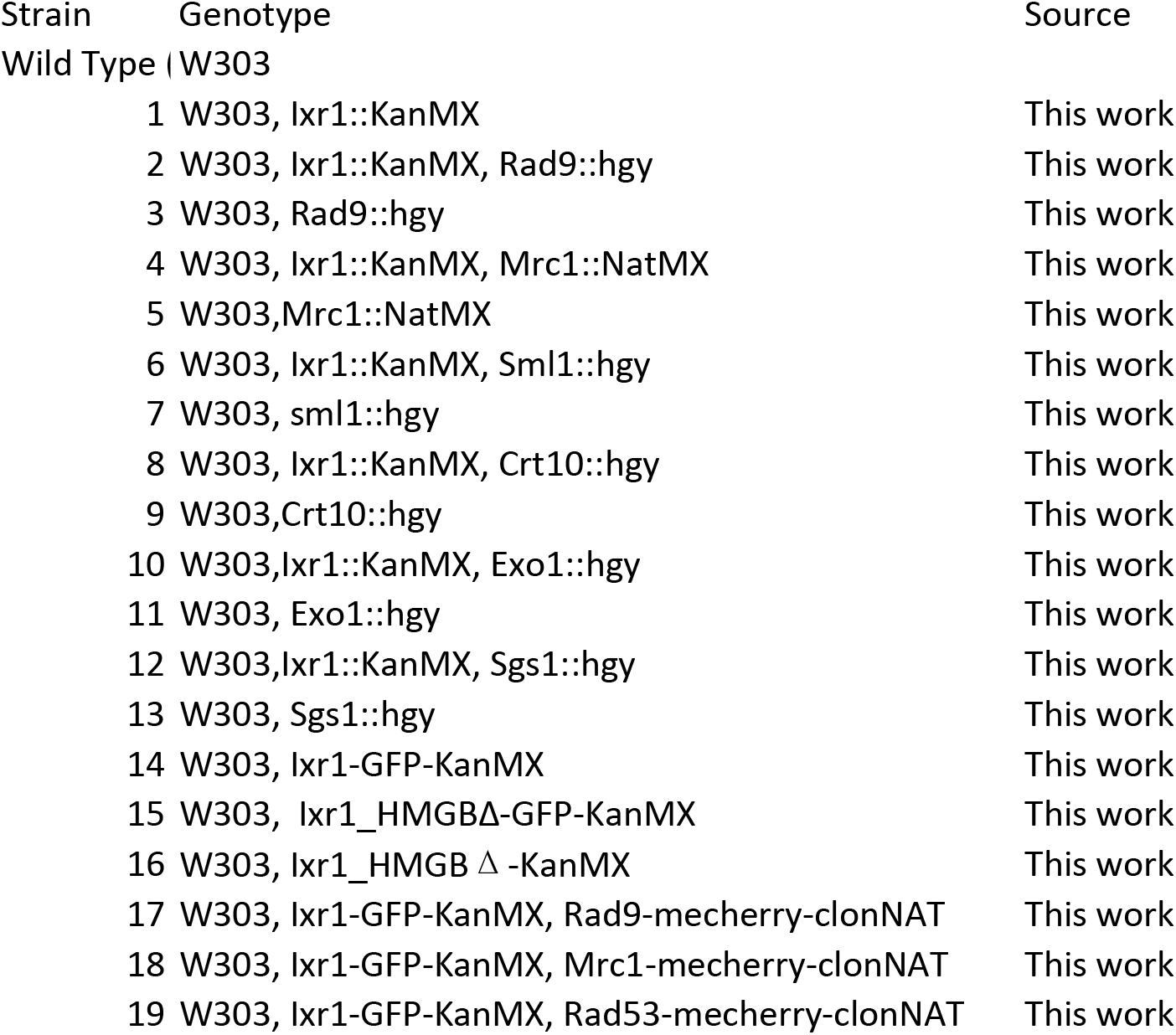

